# A Thioredoxin reductive mechanism balances the oxidative protein import pathway in the intermembrane space of mitochondria

**DOI:** 10.1101/2021.06.22.449413

**Authors:** Mauricio Cardenas-Rodriguez, Phanee Manganas, Emmanouela Kallergi, Ruairidh Edwards, Afroditi Chatzi, Erik Lacko, Kostas Tokatlidis

## Abstract

Mitochondria biogenesis crucially depends on the oxidative folding system in the mitochondrial intermembrane space. The oxidative capacity needs however to be balanced by a reductive pathway for optimal mitochondrial fitness. Here we report that the cytosolic thioredoxin machinery fulfils this critical reductive function by dual localisation in the mitochondrial intermembrane space (IMS) via an unconventional import pathway. We show that the presence of the Thioredoxin system in the IMS mediates a hitherto unknown communication between mitochondria biogenesis and the metabolic state of the cell via the cytosolic pool of NADPH. By a combination of complete in vitro reconstitution with purified components, import assays and protein interaction analysis we find that the IMS-localised thioredoxin machinery critically controls the redox state of Mia40, the key player in the MIA pathway in mitochondria thereby ensuring optimal mitochondria biogenesis. Intriguingly, we find that the IMS thioredoxin system fulfils a previously unknown role in the retrograde release of structurally destabilised proteins into the cytosol and protection against oxidative damage, both of which serve as critical mechanisms of mitochondrial surveillance and quality control.

## Introduction

Mitochondria produce most of a cell’s ATP but are also a hub for important catabolic pathways and fatty acid oxidation. Furthermore, they are involved in important processes such as the urea cycle and the synthesis of heme, cardiolipin and steroids (Scheffler, 2002). Despite possessing their own DNA, it only encodes less than 1% of the total mitochondrial proteome (about 1,500 polypeptides in mammalian cells). Thus, specialised import pathways have evolved to properly target and sort these mitochondrial proteins within the organelle’s sub-compartments (Becker *et al*, 2012, Chacinska *et al*, 2009, Chatzi *et al*, 2016, Wiedemann & Pfanner, 2017).

Among these import pathways, the **M**itochondria **I**mport and **A**ssembly (MIA) pathway accounts for the import of most of the mitochondrial **i**nter**m**embrane **s**pace (IMS) proteins (Sideris & Tokatlidis, 2010). Classical precursors of the MIA pathway contain cysteine residues in either Cx3C or Cx9C motifs (Lu & Holmgren, 2014, Lutz *et al*, 2003, Gabriel *et al*, 2007). These precursors translocate into the mitochondrion through a channel formed by the **t**ranslocase of the **o**uter **m**embrane (TOM) 40 protein in a reduced (unfolded) conformation (Stojanovski *et al*, 2012, Durigon *et al*, 2012). In the IMS, the oxidoreductase Mia40 (named CHCHD4 in humans), which possess an exposed CPC active motif, oxidises the cysteine residues on the precursor (Banci *et al*, 2009). Hence, two intramolecular disulfide bonds in the protein precursor result from this oxidation, folding and trapping the new protein in the IMS. In turn, a reduced Mia40 is re-oxidised by the protein Erv1 (**e**ssential for **r**espiration and **v**iability; Banci *et al*, 2011, Bien *et al*, 2010). An oxidative import system is not exclusive to mitochondria but is also present in the periplasm of gram-negative bacteria and the endoplasmic reticulum (ER) of mammalian cells (Denoncin & Collet 2013; Riemer *et al* 2009). However, in the bacterial periplasm and the ER a reductive system operates in the same compartment as the oxidative system to maintain the redox balance at the compartment level. Such a regulatory reductive system remains so far elusive in the mitochondrial IMS (Cardenas-Rodriguez & Tokatlidis, 2017, Herrmann & Riemer, 2014).

An analysis of the IMS proteome in *S. cerevisiae* (Vögtle *et al*, 2012) suggested the presence of thioredoxin 1 (Trx1) and the NADPH-dependent thioredoxin reductase (Trr1) in this compartment, but the import of these proteins, their potential interaction with the oxidative folding system and their function in the IMS are completely unknown. The Trx system is the key antioxidant machinery present in all cells and consists of the protein effector thioredoxin, its reductase (Trr) and NADPH as the electron source (Toledano *et al*, 2013, Draculic *et al*, 2000, Garrido & Grant, 2002, Millet *et al* 2018). Thioredoxin contains a characteristic thioredoxin (Trx) fold consisting of four central ***β***-sheets flanked by three α-helices and a conserved redox active site CxxC motif, which is located within a loop connecting a ***β***-sheet with an α-helix (Ren *et al*, 2009, Holmgren *et al*, 1975).

We need to understand the full reductive capacity of the IMS as it is crucial in the response to different types of stresses and likely in redox signalling cues in the cell. Elucidating the mechanistic aspects and redox control capacity of this dedicated Trx reductive machinery in the IMS has very broad ramifications extending to redox control of key proteins underpinning mitochondrial dynamics (Mattie et al 2018) and the balance of mtROS.

Here we show that the cytosolic thioredoxin system consisting of Trx1, and thioredoxin reductase Trr1 is dually localised in the mitochondrial IMS in addition to the cytosol in yeast *S. cerevisiae* cells. We find that import of these proteins operates via an unconventional pathway that combines several intriguing features: (i) it is independent of the key IMS import protein Mia40, (ii) it does not require ATP hydrolysis or the inner membrane potential, two known main energy sources for powering mitochondrial protein import. Moreover, the cysteine residues that are key for the redox function of the Trx system are dispensable for import, whilst prior denaturation of the protein is not prerequisite to ensure import competency. Trx1 and Trr1 are imported independently of each other, but the Trx1 localised to the IMS after import requires the IMS-localised Trr1 to remain reduced and active in the IMS. We have fully reconstituted the system with purified components and showed that the Trx machinery controls the redox state of Mia40. We further confirmed this novel Trx-Mia40 interaction also exists *in organello,* where it affects the import of proteins that depend on the MIA pathway.

Furthermore, we found that the MIA pathway is subject to control by the cytosolic NADPH, which is primarily maintained by the glucose-6-phosphate dehydrogenase and the pentose phosphate pathway. This reveals a hitherto unknown communication between mitochondria biogenesis and the metabolic state of the cell. This communication becomes critical because of the presence in the IMS of the thioredoxin system that depends on NADPH to operate. Finally, we reveal two additional functions of the Trx system that warrant further investigation in the future. First, we find that the Trx machinery assists in the retro-translocation pathway of release of structurally destabilised mitochondrial IMS proteins into the cytosol that are cleared by the cytosolic proteasomal system. Second, we provide data that support a more general role for the IMS Trx system as a crucial regulator of redox balance in this compartment protecting IMS proteins from mtROS-induced oxidative damage by carbonylation, which is a major hallmark of oxidative stress-related disorders linked to mitochondrial dysfunction, obesity and metabolic defects.

## Results

### Trx1 and Trr1 are imported into the IMS of mitochondria following an unconventional import pathway

The thiol-reductant thioredoxin proteins are evolutionarily conserved and widely distributed among species (**Figure 1A**).

**Figure 1.**
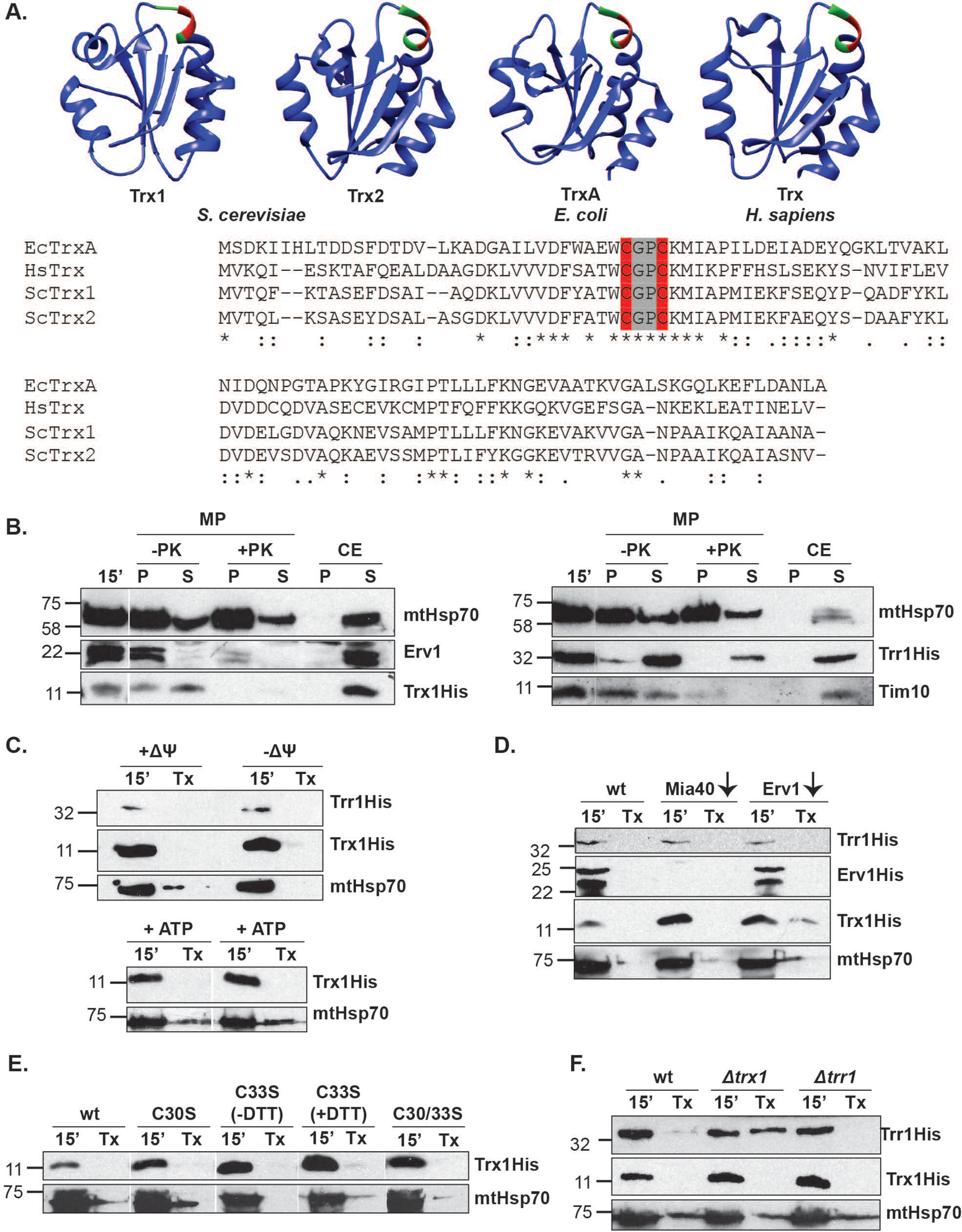
**Trx1 and Trr1 are imported into the IMS of mitochondria** A. Trx1 is highly conserved in evolution. The 3D structures of Trx1 and Trx2 from *S cerevisiae*, TrxA from *E.coli* and Trx from *H. sapiens* are shown. In the sequence alignment the strictly conserved active site CGPC is shown (in grey), with the functional Cysteines shown in red. B. Localisation of Ttrx1 and Trr1 in the IMS. Carbonate extraction (‘CE’) and mitoplasting of mitochondria (‘MP’) after the import of His-tagged recombinant Trx1 or Trr1. Samples were either treated or not with Protease K (‘+PK’ or ‘-PK’). Supernatant (‘S’) and pellet (‘P’) fractions were analysed. Anti-mtHsp70, anti-Erv1 and anti-Tim10 antibodies were used to detect the endogenous mtHSP70 (matrix-localised) and Erv1 or Tim10 (IMS-localised) as control. Lane 1 indicates the 15 min of import sample (‘15’’) as control of import. C. Import of Trx1 is independent of inner membrane potential (‘Δψ’) and ATP. ‘Tx’ is the control of Triton X100 solubilisation of mitochondria after import and treatment in the presence of externally added trypsin. D. Import of Trx1 and Trr1 is independent of Mia40 or Erv1. Import was performed in either the wild-type (wt) or Mia40-depleted (Mia40↓) or Erv1-depleted (Erv1↓) mitochondria. Detection was done using rabbit polyclonal antibodies anti-His for imported Trx1His or Trr1His, anti-Erv1 antibodies for detection of Erv1 and anti-mtHsp70 antibodies for detection of mtHsp70.’Tx’ control and 15’ of import were as in panel C. E. Import of Trx1 and is independent of its cysteine residues. Different versions of purified Trx1 as 6His-tagged proteins (wild-type wt, C30S single mutant, C33S single mutant and C30/C33S double mutant) were imported into wild-type mitochondria. Imports and controls were done as in C and D. F. Import of Trx1 and Trr1 is independent of each other. Import of pure Trx1His or Trr1His was performed into wild-type mitochondria (‘wt’), or mitochondria from a *Δtrx1* strain or from a *Δtrr1* strain as indicated on the top. Detection was with anti-His antibodies for imported Trx1His or Trr1His, and anti-mtHsp70 for endogenous mrHsp70 as control.

To study the presence of the cytosolic Trx1 and Trr1 and their unknown function in the mitochondrial IMS, we first established their import into isolated mitochondria. To this end, we purified recombinant His-tagged *S. cerevisiae* Trx1 and Trr1 from *E. coli* and imported them into wt isolated *S. cerevisiae* mitochondria. Trx1 was imported in a native state whilst Trr1 was denatured and reduced prior to import. After import, samples underwent extraction with sodium carbonate extraction and osmotic shock to establish the submitochondrial localisation of imported Trx1 and Trr1. External digestion with trypsin protease after import into mitochondria showed that both proteins (detected by anti-His antiserum in immunoblotting) have indeed crossed the outer mitochondrial membrane and are localised in a protease-protected mitochondrial localisation (‘M’ in lane 2, **Figure 1B**). Sodium carbonate extraction followed by 30-min centrifugation at 55,000g released both Trx1 and Trr1 into the supernatant fraction, showing that they are either soluble proteins or weakly associated to the inner membrane (‘CE’ treatment, lanes 7 and 8, **Figure 1B**). As a control, the endogenous IMS mitochondrial soluble proteins Erv1 and Tim10 and mtHsp70 (a matrix protein) were also detected by specific antisera to be quantitatively released in the supernatant. In the osmotic shock treatment, the mitochondrial outer membrane was disrupted by hypotonic swelling yielding mitoplasts (MP). A further centrifugation step separates the MP (pellet) from the IMS fraction (supernatant) (Lanes 3-6 **Figure 1B**). As a control, both Erv1 and Tim10 proteins were degraded by PK in the mitoplast fraction, in contrast to the matrix-localised mtHsp70 which remained largely intact. Both Trx1 and Trr1 were present mostly in the supernatant, whilst addition of protease K (PK) during the generation of mitoplasts substantially degraded both proteins, confirming their localisation in the IMS (compare lanes 3 and 4 to 5 and 6, **Figure 1B**).

Next, to get insight into the import mechanism for both proteins, we examined the requirement for ether ATP or the mitochondrial electrochemical potential of the inner membrane, as these are the two major energy sources that power protein import into mitochondria. Depletion of ATP was achieved by treating the mitochondria with oligomycin and apyrase and removing the externally supplied ATP during the import process (Terziyska *et al* 2007), whilst depletion of the inner membrane potential was achieved by treating them with valinomycin. Depletion of ATP or disruption of the mitochondrial inner membrane potential (Δψ) did not affect the import of Trx1 and Trr1 into isolated wt mitochondria (**Figure 1C)** showing that, these energy sources are not required for the import of Trx1 and Trr1 into mitochondria.

The MIA machinery provides the main import pathway for the IMS compartment, particularly for cysteine-containing proteins. In this case, it is thought that the formation of internal disulfide bonds facilitated by the MIA machinery provides the energy to retain the proteins that follow this pathway in a folded state in the IMS. We therefore tested whether the absence of the main components of the pathway, Mia40 and Erv1, impacts the import of Trx1 and Trr1. As a control, the import of Erv1 (an import substrate for Mia40) was abolished in the Mia40-depleted mitochondria. Intriguingly, import of Trx1 and Trr1 into isolated mitochondria from yeast cells with conditionally-depleted expression of either Mia40 (GalMia40) or Erv1 (GalErv1) showed no decrease in the import levels of the Trx proteins compared to import in wt mitochondria (**Figure 1D**). In fact, the import of Trx1 in the Mia40 and Erv1-depleted mitochondria seems to be increased. This is an interesting observation which would require further work, beyond the scope of this paper, to clarify in the future.

Cysteine-containing proteins of the IMS are usually dependent on these cysteines for import. For example, the classic substrates of the MIA pathway need to form mixed disulfide intermediates with Mia40 through their cysteine residues to be imported. We tested whether the cysteines of Trx1 played a role in the import pathway by generating cysteine to serine mutants of Trx1 (C30S, C33S and C30/33S) and importing these into wt mitochondria. All three cysteine to serine mutants displayed no import defects, showing that the cysteine residues do not play a role in the import of Trx1 into mitochondria (**Figure 1E**).

Given that the import of Trx1 and Trr1 was found to be independent of the MIA pathway (**Figure 1D**) and did not require the inner membrane potential or ATP (**Figure 1C**), we investigated whether the presence of the one protein in the IMS guided the import of the other. Such a mechanism would be akin to the import pathway of other proteins dually localised in the cytosol and the IMS and partner each other in the IMS. One such example is the import of Sod1 which is dependent on the prior import of its copper chaperone Ccs1 (Field *et al*, 2003, Kawamata & Manfredi, 2008). However, the situation for Trx1 and Trr1 proved to be different. Import of Trx1 and Trr1 into isolated mitochondria from yeast mutants lacking Trx1 (*ΔTrx1*) or Trr1 (*ΔTrr1*) showed no dependence at all on the presence of either Trx1 or Trr1 (**Figure 1F**).

Taken together, the above data show that the cytosolic isoforms of the thioredoxin system, Trx1 and Trr1, dually localise to the mitochondrial IMS in addition to their hitherto known presence in the cytosol. Furthermore, the import of these proteins in the IMS defines an unconventional pathway which is distinct from the main known import pathways to the IMS and is independent from the presence of Trx1 and Trr1 themselves in the IMS.

### Trx1 stays reduced in the IMS in a Trr1-dependent manner and binds to Mia40 *in vitro* and in organello

In the cytosol, Trx1 is reduced by thioredoxin reductase (Trr1) in the presence of NADPH which serves as the electron donor for this system to operate (Toledano *et al*, 2013). Keeping Trx1 in a reduced state by Trr1 is a critical requirement for the function of Trx1. To confirm whether this mechanism is active for the pool of Trx1 and Trr1 proteins that localise in the mitochondrial IMS, we imported the recombinant His-tagged Trx1 into either wt mitochondria or mitochondria isolated from a yeast strain that lacks Trr1 (*Δtrr1*) followed by an alkylation shift assay to assess the dependence of the redox state of the IMS-localised Trx1on Trr1. The alkylating molecule AMS (4-acetamido-4’-maleimidylstilbene-2,2’-disulfonic acid) binds to reduced sulfhydryl groups (-SH), but not to oxidised S-S groups. This modification causes a visible gel shift in SDS-PAGE of 0.5 kDa for every AMS molecule bound to one sulfhydryl group. By performing this assay for Trx1 after import in mitochondria, we saw that Trx1 is maintained in a reduced state when imported into wt mitochondria (which contain endogenous Trr1 in the IMS) but, is instead oxidised when imported into mitochondria lacking Trr1 from a *Δtrr1* strain (**Figure 2A**).

**Figure 2.**
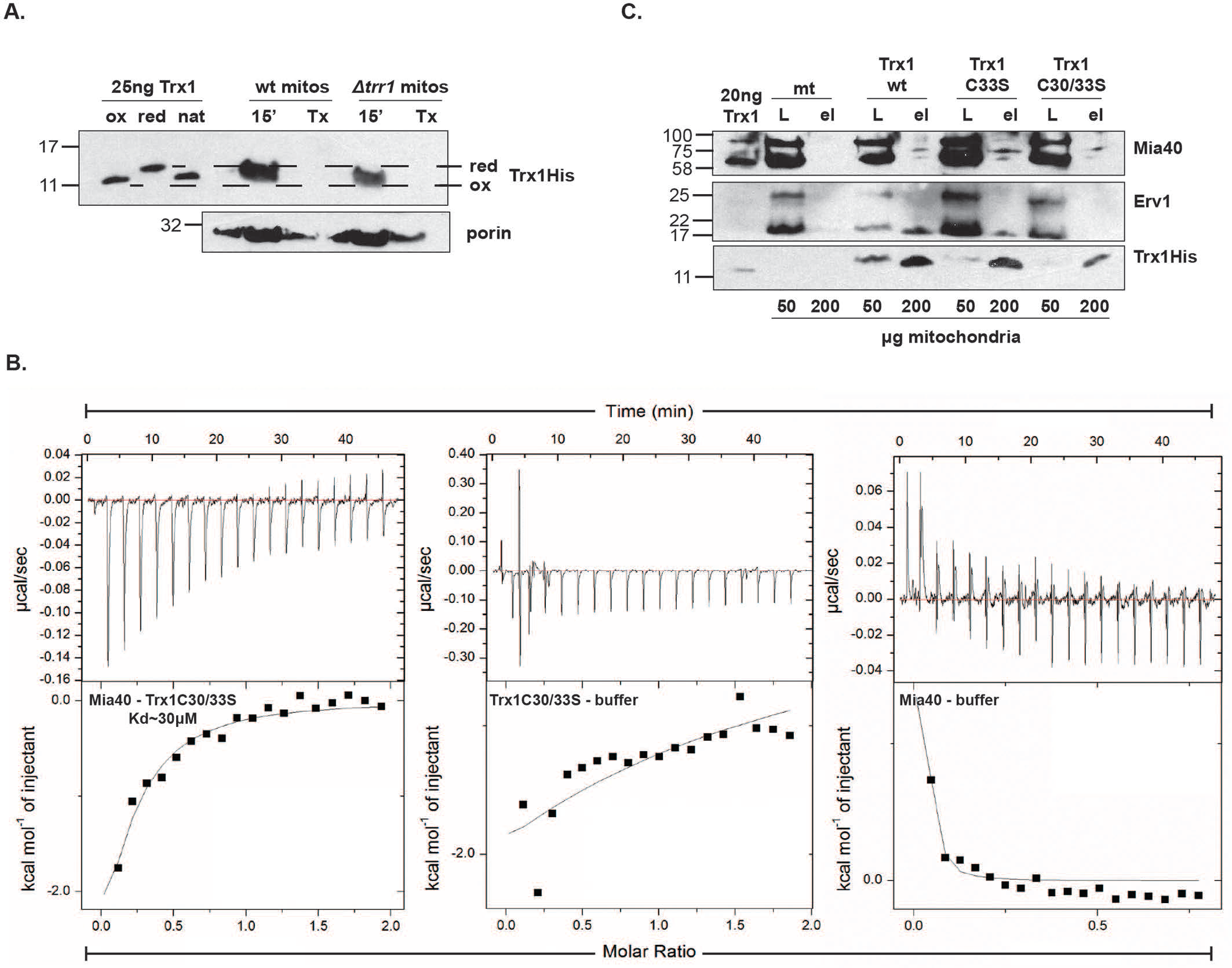
**The Trx system interacts with Mia40 *in vitro* and *in organello*** A. Reduction of Trx1 within mitochondria depends on the presence of Trr1. In the first three lanes the pure Trx1 was either completely oxidised (‘ox’) or completely reduced (‘red’) or loaded as purified without further treatment (‘nat’). The dotted lines were drawn to help distinguish the different mobility of the reduced and oxidised forms. Trx1His was imported either in wild-type mitochondria (that contain Trr1 protein in the IMS) or in mitochondria from a *Δtrr1* train (lacking Trr1). AMS thiol-trapping was used to distinguish the reduced (AMS bound and hence showing a larger band shift) and oxidised forms (AMS not bound and migrating faster as a smaller protein). Porin was used as endogenous protein control. ‘Tx’ is the control of Triton X100 solubilisation of mitochondria after import in the presence of trypsin to show that Trx1 is indeed imported. B. Trx1 and Mia40 interact *in vitro* as shown by isothermal titration calorimetry. The Trx1 double mutant C30/33S was used to avoid thiol disulfide exchange with Mia40 and only detect non-covalent binding interactions (left panel). As control, the Trx1C30/33S against buffer only (middle panel) and Mia40 against buffer only (right panel) are shown. C. Trx1 and Mia40 interact *in organello*. Different versions (wt, the C33S single mutant which is a thiol-trapping mutant, and the double mutant C30/33S) of purified Trx1His were imported into wt mitochondria (indicated on the top of the gel), followed by Ni-NTA binding, SDS-PAGE and analysis by immunoblotting. Anti-His antibodies were used to detect Trx1, and anti-Mia40 or anti-Erv1 rabbit polyclonal antibodies were used to detect Mia40 and Erv1 respectively. ‘L’: total loading material; ‘el’: eluate after binding to the Ni-NTA beads. ‘mt’: mitochondria without any TrxHis pre-imported. The amount of mitochondria used is indicated in the bottom of the gel, and the molecular weight markers to the left of the gel.

Next, we wanted to investigate whether the Trx system interacts with Mia40, given the fact that the function of the essential Mia40 protein is critically controlled by its redox state (Milenkovic *et al*, 2009, Sideris *et al*, 2009, Sideris & Tokatlidis, 2010). We addressed this question in experiments done both *in vitro* with purified proteins and *in organello* with isolated mitochondria. First, we assessed the binding of purified Mia40 and Trx1 using isothermal titration calorimetry, a method that has been used in the past to measure the thermodynamic affinity of Mia40 to its substrates and to Erv1(Sideris *et al*, 2009 and Lionaki *et al*, 2010). For these experiments, and all subsequent experiments for *in vitro* binding of Mia40, we used the *S. cerevisiae* Mia40 version that is devoid of the N-terminal 290 residues which are dispensable for function (Naoé *et al*, 2004, Sideris *et al* 2009). This core domain of Mia40 (about 15 kDa, Mia40Δ290 His) retains full function and is similar to the smaller human Mia40 which does not contain the N-terminal extension that is only present in the *S. cerevisiae* Mia40 protein (Naoé *et al*, 2004; Chacinska *et al*, 2004). Using this soluble functional domain of Mia40 we found that it binds to Trx1 with a strong affinity and a Kd of 30 μM. (**Figure 2B**) This is comparable to the affinity of Mia40 to Erv1 (25μM) (Lionaki *et al*, 2010) that re-oxidises Mia40, and is weaker than the binding of Mia40 to its substrates like Tim10 (3.3μM) (Sideris *et al,* 2009)

We further assessed the *in vitro* interaction between the Trx1 and Mia40 using a chemical crosslinking approach with the cross-linker glutaraldehyde (GA). We identified several bands at 12kDa, 27kDa, 35kDa and 52kDa representing monomers and likely oligomers within themselves when cross-linking each individual protein with GA (Trx1-Trx1 or Mia40-Mia40) (**Supplementary Figure 1**). However, when Trx1 and Mia40 were incubated together in the presence of GA, a new band at 30kDa appeared in addition to all the other bands that were visible when the individual proteins were crosslinked on their own. This likely represents a Mia40-Trx1 adduct. Similar results were obtained with Trx2 which is almost in sequence to Trx1. To ascertain whether the crosslinking Trx1-Mia40 interaction reflected a protein-protein interaction independent of the cysteines of Trx1, we performed the same cross-linking experiment using the C30/33S Trx1 redox-inactive version. The same 30 kDa Mia40-Trx1 adduct band was still present, suggesting the cysteines are not strictly required for the non-covalent protein-protein interaction. (**Supplementary Figure 1**). We also observed that in addition to the oligomeric species of the Trx1 sample, additional higher molecular weight bands are present, probably reflecting subtle structural changes of C30/C33 mutant compared to the wt Trx1.

To address whether the interaction of Trx1 with Mia40 also occurs *in organello*, we performed pulldown experiments using isolated mitochondria. Trx1His wt, Trx1His C33S and Trx1His C30/33S were imported into wt mitochondria, which were then solubilised with 0.5% n-dodecyl-maltoside. The solubilisation supernatant was incubated with equilibrated Ni-NTA beads. We can see that Trx1His could pull down Mia40. The relatively low levels of Mia40 pulled down is likely due to the fact that Trx1 interacts with several proteins in the IMS and Mia40 itself is also binding several of its import substrates allowing a small fraction to be pulled down by Trx1. The cysteine trap mutant (Trx1His C33S) was able to pull down higher amounts of Mia40 than the Trx1His wt, indicating a thiol-exchange interaction between the two proteins. On the other hand, the levels of Erv1 pulled down by Trx1 are extremely low and suggest that there is a much stronger interaction between Trx1 and Mia40. The non-imported mitochondria (mt) and the Trx1His C30/33S-imported mitochondria were used as controls for the detection of any background signals. These results are in agreement with the *in vitro* binding results, and show that Trx1 and Mia40 interact in mitochondria (**Figure 2C**).

### Reconstitution of the Mia40 reduction by the Trx1/Trr1 system using purified components

To investigate whether the physical binding observed between Trx1 and Mia40 in **Figure 2** is functional, *i.e*. whether Trx1 reduces Mia40, we reconstituted the entire system *in vitro* with purified components using Mia40 as a substrate on the one hand and the Trx1 and Trr1 as the reductive machinery on the other with NADPH as the physiological source for providing reductive equivalents to the Trx system. The redox state of Mia40 was monitored in a gel-shift assay followed by AMS-alkylation to detect a redox shift in Mia40 from its oxidised to its reduced state.

We first assessed whether this *in vitro* reconstituted Trx system was functional in obtaining reduced (and therefore active) Trx1 after addition of both Trr1 and NADPH (**Figure 3A**). Lanes 1 and 2 show the fully oxidised (treated with H_2_O_2_) or fully reduced (treated with DTT) purified Trx1 (with an obvious shift in migration between the two forms), while lanes 3 and 4 contain the purified Trr1. Trr1 was treated with 5mM NADPH for 30min prior to performing the reaction with Trx1. The sample in lane 4, where Trr1 was treated with DTT, was added as a control to show how Trr1 migrates in the gel when reduced. The interaction between Trx1 and Trr1, where Trr1 together with NADPH was added to oxidised Trx1, is shown in lanes 5-9 (kinetics from 30 sec to 10 minutes). We observe that the Trx1 becomes fully reduced (‘Trx1-AMS_2_’) upon addition of Trr1/NADPH, already within the first 30 seconds of the reaction.

**Figure 3.**
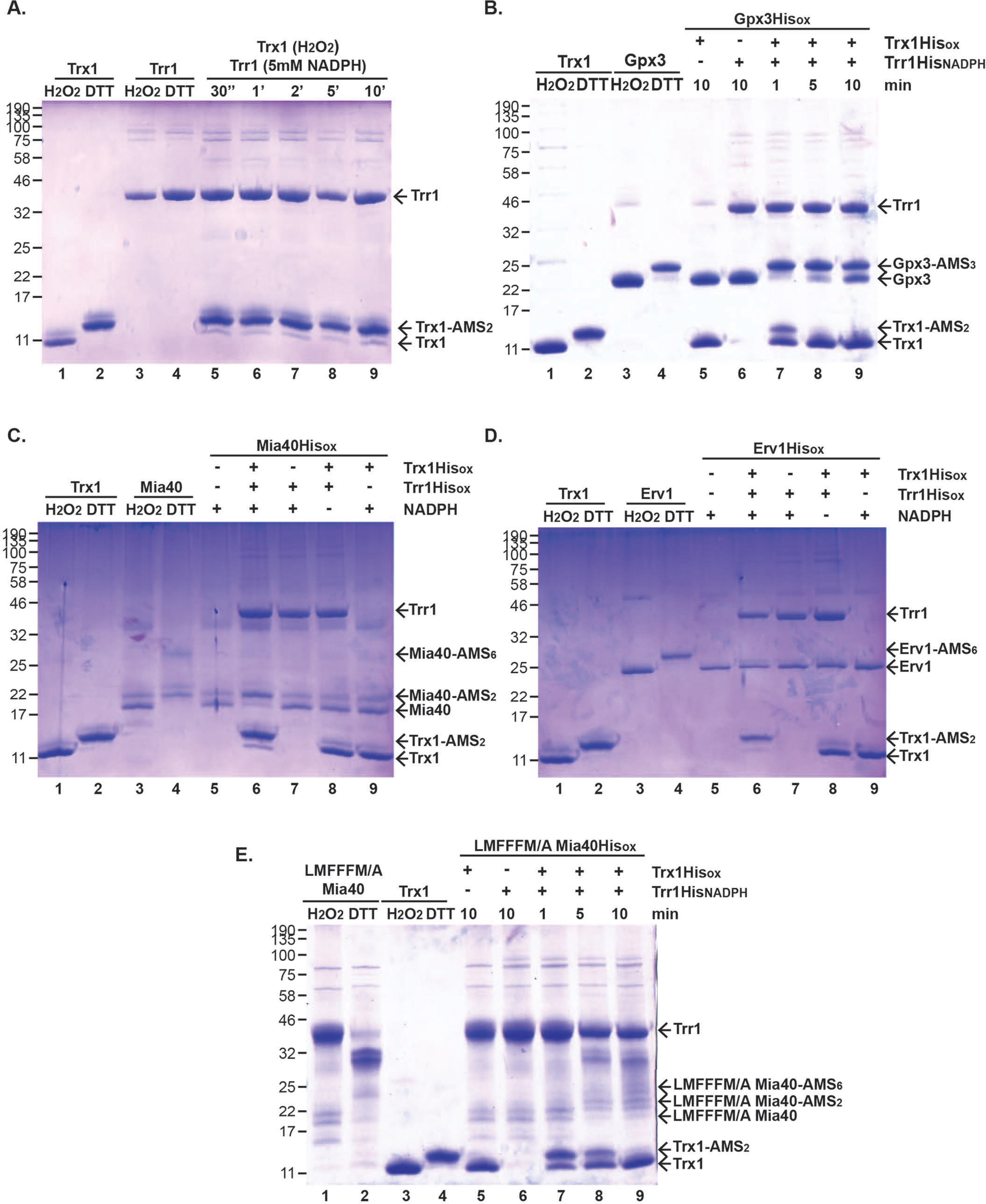
**Complete *in vitro* reconstitution of the reduction of Mia40 by the Trx system using purified components** A. Trx1 is fully reduced by Trr1 and NADPH in a minimal *in vitro* system using purified proteins. B. Lanes 1-2: Trx1 alone; Lanes 3-4: Trr1 alone. Fully oxidised Trx1 and Trr1 were treated with H_2_O_2_ (‘H_2_O_2_’, lanes 1 and 3 respectively), whilst fully reduced were treated with DTT (lanes 2 and 4). Lanes 5-9: Trx1, Trr1 and NADPH were added (see text for details). The fully reduced Trx1 binds two molecules of AMS and indicated as ‘Trx1-AMS_2_’. Incubation times for the Trx1/Trr1/NADPH mixture were from 30 sec to 10 min as indicated (lanes 5-9). B. The fully *in vitro* reconstituted Trx system can reduce the control protein Gpx3. Lanes 1-2: Trx1 alone (as in A); Lanes 3-4: Gpx3 alone (oxidised by H2O2 in lane 3, or reduced by DTT in lane 4); Lanes 5-9: Components of the Trx machinery added (as indicated) to oxidised Gpx3; Lane 5: only TrxHis oxidised added (as control); Lane 6: only Trr1His and NADPH added; Lanes 7-9: the full Trx machinery i.e Trx1 and Trr1 and NADPH added. C. The fully *in vitro* reconstituted Trx system can reduce Mia40. The experiment and analysis were performed in an identical manner as in panel B. Lanes 1-2: Trx1 alone; Lanes 3-4: Mia40 alone; Lanes 5-9: MiaHis-oxidised was incubated with different combinations of components of the Trx machinery as indicated. D. The Trx system does not interact with Erv1, showing a specificity to Mia40 in the oxidative folding machinery of the IMS. The experiment and analysis were done exactly as in C, but using Erv1 instead of Mia40. E. The hydrophobic cleft of Mia40 is not critical for the interaction with the Trx system. The experiment and analysis were done exactly as in C, but using the LMFFFM/A mutant in the substrate binding cleft of Mia40 instead of wild-type Mia40.

Next, we wanted to confirm that this reconstituted Trx system can indeed work with a known substrate for Trx1 before embarking on the assay using Mia40. As a known, well-studied substrate for Trx1 we used the thiol peroxidase Gpx3, which is reduced by the thioredoxin system in the yeast cytosol (Delauney *et al*, 2002). (**Figure 3B**). Lanes 1 and 2 show the fully oxidised (treated with H_2_O_2_) or fully reduced (treated with DTT) purified Trx1, while lanes 3 and 4 contain the purified Gpx3. For both proteins the respective reduced forms are modified by AMS (Trx1-AMS_2_ or Gpx3-AMS_3_) resulting in a slower mobility compared to the AMS-unmodified oxidised forms. Addition of oxidised Trx1 to oxidised Gpx3 does not reduce Gpx3 at all, as expected (lane 5, **Figure 3B**). Additionally, addition of Trr1/NDPH but in the absence of Trx1 does not reduce the oxidised Gpx3 either (lane 6, **Figure 3B**). By contrast, addition of Trx1 together with Trr1 and NADPH (which represents the full reductive system) results in reduction of the oxidised Gpx3 (lanes 7-9). Looking at the kinetics of this reaction, we observe that already within one minute Gpx3 is almost fully reduced and concomitantly Trx1 is becomes oxidised (lane 7); as the reaction progresses (to 5 and 10 minutes in lanes 8 and 9), Trx1 is getting quantitatively oxidised as a substantial part of Gpx3 is maintained in a reduced state.

Having established this functional reconstitution of the Trx system with a known substrate, we tested the reduction of Mia40 (**Figure 3C**). Indeed, we find that that the Trx system efficiently reduces Mia40 in the *in vitro* reconstitution assay. Mia40 contains three cysteine pairs, the active site CPC pair which readily undergoes reduction and oxidation, and the two structural cysteine pairs connecting the twin CX9C motifs which are highly resistant to reduction (Chacinksa *et al* 2004; Grumbdt *et al*, 2007; Banci *et al,* 2009). Lanes 1 and 2 in **Figure 3C** contain the oxidised and reduced forms of Trx1 while lanes 3 and 4 contain the oxidised and reduced form of Mia40. In lane 4 we note two main forms of AMS-labelled Mia40; the fully reduced Mia40-AMS_6_ with 6 AMS molecules alkylating all six cysteines of Mia40, and the partially reduced Mia40-AMS_2_, with 2 AMS molecules bound to the more readily reducible cysteines of the CPC motif. Different combinations of Trx1, Trr1 and NADPH were then added to oxidised Mia40 to elucidate how it can be reduced by the Trx system. Interestingly, we observed a substantial, almost quantitative reduction in the active site CPC motif (Mia40-AMS_2_) only when Trx1 was added together with Trr1 and NADPH (lane 6, **Figure 3C**), which constitutes the full Trx1 reductive machinery. By contrast, NADPH alone (lane 5), Trr1 with NADPH but without Trx1 (lane 7), Trx1 and Trr1 but without NADPH (lane 8), or Trx1 with NADPH but without Trr1 (lane 9) were all insufficient to substantially reduce Mia40 in its CPC motif.

These results therefore support that the reduction of the active site CPC motif of Mia40 by the Trx1/Trr1/NADPH system can be fully reconstituted in an *in vitro* minimal system using purified components.

Is the thioredoxin-mediated reduction specific for Mia40? To address this, we used the other critical component of the mitochondrial oxidative folding pathway the FAD-oxidoreductase Erv1 as a putative substrate in the reconstitution assay. As can be seen in **Figure 3E**, in contrast to Mia40, Erv1 cannot get reduced by the thioredoxin system. Erv1 can be fully reduced by DTT with all 6 of its cysteines alkylated by AMS (Erv1-AMS_6_) as can be seen in lane 4, whereas the fully oxidised form of Erv1 (with H_2_O_2_) is not modified by AMS (lane 3 **Figure 3D**). Interestingly, none of the conditions used in the reconstitution assay (lanes 5-9 **Figure 3D)** could reduce substantially Erv1, including the optimal combination of Trx1/Trr1/NAPDH (lane 6, **Figure 3D**) which only reduces Erv1 at extremely low levels, in contrast to the situation where the same conditions could however reduce Mia40 quantitatively (lane 6, **Figure 3C**). This data therefore supports a mechanism whereby the Trx1/Trr1/NADPH reductive machinery very efficiently specifically reduces Mia40.

The common determinant for all the so far known functional interactions of Mia40 (interaction with its substrates and with Erv1) is its hydrophobic cleft; this ensures (i) its chaperone binding to the substrates (Sideris *et al,* 2009, Banci *et al,* 2009) and (ii) the capacity of Erv1 to bind to Mia40 and re-oxidise it by mimicking the substrate (Banci *et al*, 2011). It was therefore interesting to investigate whether this crucial hydrophobic domain also underpinned the reduction of Mia40 by Trx1. To address this, we used a mutant version of Mia40 in the reconstituted system, where the amino acid residues of the hydrophobic cleft were replaced by alanine residues (LMFFFM/A, Banci *et al*, 2009)). This mutant was previously shown to lose the capacity to interact with either the Mia40 substrates or with Erv1 (Chatzi *et al,* 2013). Intriguingly, reduction of this Mia40 mutant by Trx1 remained unaffected, suggesting that the binding of Trx1 to Mia40 is independent of its hydrophobic cleft and therefore very different from the binding of substrates or Erv1 in structural terms. (**Figure 3E**). This represents a potentially new interaction mechanism for Mia40, unrelated to the way the substrates bind or the substrate mimicry mechanism of recycling by Erv1 (Banci *et al,* 2011)

### The redox balance of Mia40 in the IMS is under the control of the cytosolic pool of NADPH and affects its key function in the MIA import pathway

Mia40 oxidises free sulfhydryl groups within its classical protein substrates, thereby catalysing formation of internal disulfide bonds in these proteins that stabilise their structure. This mechanism is key to retaining these Mia40 substrate proteins in the mitochondrial IMS (Banci *et al*, 2011, Banci *et al*, 2009Bien *et al*, 2010, Chacinska *et al*, 2004). It has recently been shown in human cells that Mia40 is in a semi-oxidised state (Erdogan *et al*, 2018). Therefore, maintaining a balance between the oxidised and reduced state of Mia40 is critical for its optimal function. Since the mitochondrial IMS pool of Trx1 interacts with Mia40 and is able to reduce it in *in vitro* reconstitution assays, we wondered whether Trx1 physiologically impacts the redox state of Mia40 thereby affecting its import capacity. We used isolated mitochondria from a yeast strain that lacks the yeast enzyme glucose 6-phosphate dehydrogenase (*Δzwf1*, noted for simplicity as *ΔG6PD* in the present work), the main source of NADPH, to determine the redox state of Mia40, assess protein levels and perform import experiments. First, we determined that this *ΔG6PD* yeast strain is sensitive to respiratory conditions. We performed spot test analysis where we grew serial dilutions of yeast under non-respiratory (YPD) and respiratory (YPLac) conditions at permissive (30℃) and non-permissive (37℃) temperature. We compared the ΔG6PD yeast with yeast lacking the main reductant effectors: both cytosolic versions of Trx (*Δtrx1Δtrx2*), Trr1 (*Δtrr1*) or the Gsh1 enzyme (*Δgsh1*), which catalyses the first step in GSH synthesis. No difference was observed between the yeast mutants and the wt under non-respiratory growing conditions at either temperature (Figure 3A). However, when the yeast cells were grown in the presence of a respiratory carbon source, clear growth defects were observed for both *ΔTrx1* and *ΔGsh1* strains, and an even more pronounced defect was observed for the *ΔG6PD* mutant (**Figure 4A**).

**Figure 4.**
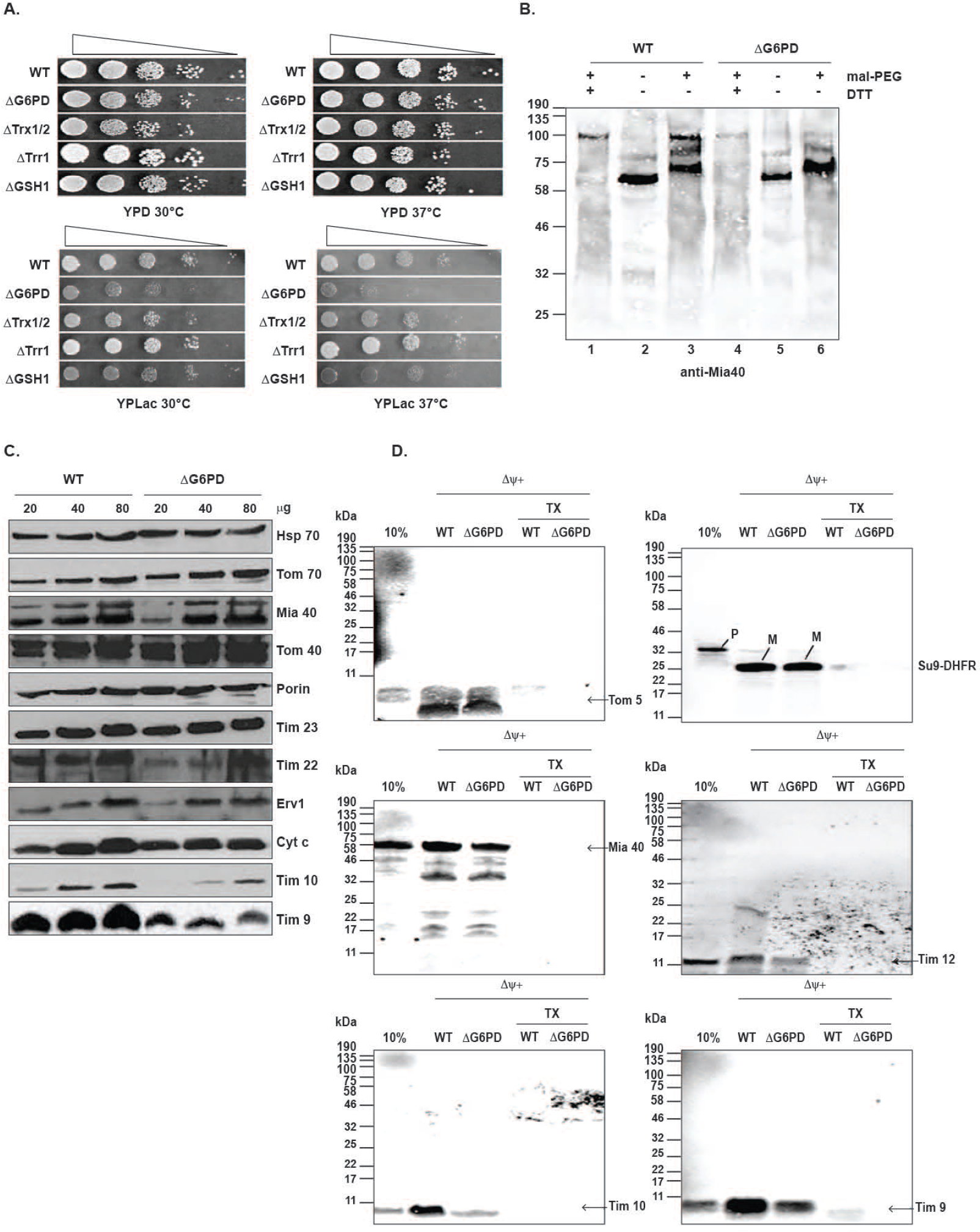
**The redox state of Mia40 is influenced by cytosolic NADPH in a G6PD mutant thereby compromising its import capacity.** A. Growth of mutants of the main reducing pathways is impaired under respiratory conditions (lactate media, YPlac) compared to fermentative conditions (glucoce media, YPD). The *Δzwf1* (null mutant in the *S. cerevisiae* Glucose-6-phosphate dehydrogenase; indicated as ΔG6PD) shows the most pronounced defect. B. Redox state of Mia40 in Δ*zwf1* is mainly oxidised as opposed to the wild-type cells. Mitochondria were purified from the indicated yeast strains and total mitochondrial extracts were used for thiol trapping with mal-PEG (which adds 5,000 Dalton for every free thiol group) as indicated. Samples were analysed on SDS-PAGE followed by immunoblotting with anti-Mia40 rabbit polyclonal antibodies. C. Steady state levels analysis with immunoblotting of mitochondrial proteins; Levels of proteins that are import substrates of Mia40 (Tim9, Tim10, Tim22) are clearly reduced in *Δ zwf1* mitochondria (indicated as ΔG6PD), while other mitochondrial proteins are unaffected. D. Import of mitochondrial precursors into WT and ΔG6PD mitochondria shows a specific defect for MIA substrates (Tim9, Tim10, Tim12) but not for proteins imported via other pathways (Tom5, Su9DHFR, Mia40). Import assays were done using 35S-labelled precursors, followed by autoradiography. ‘TX’ indicates a control where mitochondria were completely solubilised by Triton X-100 followed by trypsin digestion. All imports were done in the presence of a functional IM potential (Δψ+).

The reducing power of NADPH feeds both of the two main reducing systems of the cell: the Trx and glutaredoxin (Grx) systems (Toledano *et al*, 2013). To pinpoint any effect of a compromised NADPH reductive cellular capacity on Mia40, we isolated mitochondria from the *ΔG6PD* yeast cells and looked at the redox state of Mia40 using an alkylation shift assay as previously described but utilising mal-PEG (which adds 5kDa for each free sulfhydryl group) as an alkylating agent. We found that in wt cells Mia40 is present in a mixed population of reduced and oxidised states (lanes 1-3, **Figure 4B**). By contrast, in the *ΔG6PD* mitochondria the redox balance is lost and Mia40 is mainly shifted to its oxidised form (lanes 4-6, **Figure 4B**).

Does this altered redox balance of Mia40 affect other mitochondrial proteins? To address this, we examined the levels of mitochondrial proteins in *ΔG6PD* mitochondria in comparison with wt mitochondria by immunoblotting. Interestingly, the levels of IMS proteins that depend on Mia40 for their import such as Tim9 and Tim10 were found to be substantially decreased in *ΔG6PD* mitochondria, while proteins of the other sub-mitochondrial compartments (outer membrane, inner membrane and matrix) that are imported via Mia40-unrelated pathways are unaffected. The slightly lower levels of Tim22 in the *ΔG6PD* mitochondria could be an indirect effect attributed to the lower levels of Tim9 and Tim10 which control the import of Tim22 in the carrier pathway (Sirrenberg *et al*, 1998; Koehler *et al*, 1998, Wrobel *et al*, 2013)). (**Figure 4C**).

To elucidate directly whether the observed lower protein levels were due to compromised import as a consequence of the altered redox balance of Mia40 in *ΔG6PD* mitochondria, we performed import experiments. Radiolabelled protein precursors from the four different mitochondrial compartments were presented to isolated wt or *ΔG6PD* mitochondria and the non-imported material was cleaved with proteinase K (PK). The import of Tom5, Su9-DHFR, Mia40, was not affected, whereas the import of Tim12, Tim10, Tim9 was decreased in *ΔG6PD* mitochondria (**Figure 4D**).

These results show that the absence of cytosolic NADPH specifically results in loss of redox balance in the mitochondrial IMS specifically through Mia40. Such loss of redox equilibrium for Mia40 impacts on its capacity to function properly as a crucial import component in the MIA pathway.

### Targeting of the Trx system in the mitochondrial IMS restores the redox state of Mia40 and protein import via the MIA pathway

The fact that the lack of NADPH in the cytosol is communicated to the IMS of mitochondria is important for ensuring a proper redox balance in this mitochondrial sub-compartment. Previously, it has been suggested that reduced GSH in the IMS is important as a reductant (Bien *et al*, 2010; Kojer *et al*, 2012), but the import of glutaredoxin in the IMS which would control GSH remains elusive. Given our data that the Trx system can be imported to the IMS, can bind to Mia40 and reduce it *in vitro*, we reasoned that supplying the ΔG6PD mitochondria, where a mostly oxidised form of Mia40 is present, with the Trx system, should restore a physiological redox balance for Mia40. To address this, we imported Trx1 and Trr1 into ΔG6PD mitochondria in the presence of

NADPH (**Figure 5A**), and subsequently analysed the redox state of Mia40 using mal-PEG as shown in **Figure 5B**. This resulted in an almost complete restoration of the redox balance of Mia40 to levels that are identical to wt (**Figure 5B**, see lane 3 for wt mitochondria, compared to lane 6 for *ΔG6PD* mitochondria without any addition of the Trx system, and lane 8 where the full Trx1/Trr1/NADPH was added to *ΔG6PD* mitochondria).

**Figure 5.**
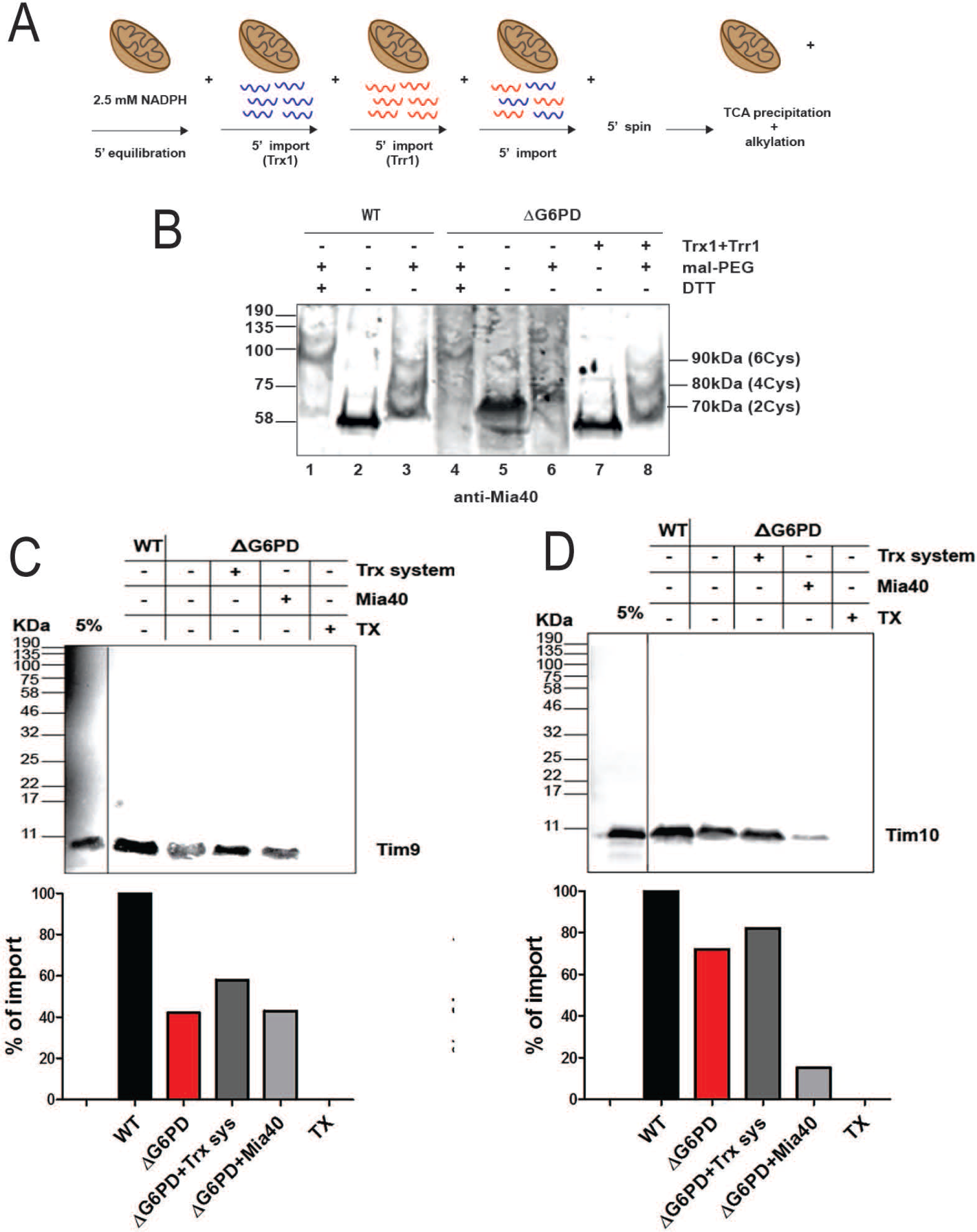
**The addition of the Trx system in the mitochondrial IMS restores the redox state of Mia40 and its import capacity.** A. Schematic representation of the step-wise addition method of the full Trx system in mitochondria. B. Pre-import of the full Trx system into *Δ zwf1* mitochondria (ΔG6PD) restores the redox state of Mia40 to WT levels. Lanes 1-3: wild-type mitochondria; Lanes 4-8: ΔG6PD mitochondria; Lanes 4-6: no addition of the Trx system; Lanes 7-8: addition of the full Trx system. The different forms of Mia40 with 2, 4 or 6 Cys in a free thiol state (and hence bound to mal-PEG) are indicated. Analysis was done similarly to Figure 4B. C. Import of the Mia40 substrate Tim9 in ΔG6PD mitochondria improves after pre-import of the Trx system. 35S-labelled Tim9 precursor was imported into pure wild-type or ΔG6PD mitochondria, followed by SDS-PAGE and autoradiography. In the first lanes of each gel 5% of the total amount of the in vitro translation mix used for import was loaded as a control. Quantification shows that addition of the Trx system (‘Trx sys’, dark grey column) is increased compared to import in ΔG6PD mitochondria without the addition of the Trx system (red column), or when the ΔG6PD mitochondria were supplemented with Mia40 alone (light grey column). D. Same as C but for import of the Mia40 substrate Tim10.

Next, we tested whether this restored redox balance of Mia40 in the ΔG6PD mitochondria by the presence of the Trx system in the IMS and the addition of NADPH also improved the Mia40 import capacity. To address this, we established a stepwise import assay where we added into *ΔG6PD* mitochondria first NADPH, then purified Trx1 and Trr1 and finally a 35S-labeled import substrate (**Figure 5A and 5C**). In these experiments the addition of the full Trx system in the IMS together with NADPH as the reductive source indeed improved the import of Tim9 and Tim10 (two well-known substrates of Mia40) (**Figure 5C**). Addition of Mia40 to the ΔG6PD mitochondria did not improve the import, arguing for the necessity of the Trx in the IMS to balance properly the redox state of Mia40. Taken together the data show that the targeting of the complete Trx1/Trr1 reductive system in the IMS with NADPH as the reductive source can restore the redox balance of Mia40, ensuring thus its optimal function in the mitochondrial IMS.

We then wanted to address whether supplementation of the IMS in mitochondria from a *Δtrx1/2* strain with the full Trx1/Trr1/NADPH system directly improved import of MIA substrates. To this end, we used Tim9 as a model MIA import substrate. **Figure 6** shows that the import of Tim9 in *Δtrx1/2* mitochondria is clearly affected. **Figure 6A** shows the SDS-PAGE analysis after import, **Figure 6B** shows the BN-PAGE analysis to monitor the assembly of imported Tim9 where we see a very strong effect, and **Figure 6C** shows the quantification of imported Tim9, with error bars from three independent experiments. The import in *Δtrx1/2* mitochondria is lowered by about 50% compared to the wt. When we supplemented the *Δtrx1/2* mitochondria with the full Trx system (**Figure 6D**, lanes 3 and 4 ‘+Trx system’), we see a noticeable improvement in the import of Tim9 (analysed at 10 and 30 min of import) compared to the import without supplementation with the Trx system (**Figure 6D**, lanes 1 and 2 ‘-Trx system’). This result is mirrored by an improvement in the assembly of Tim9 analysed by BN-PAGE (**Figure 6E**). The quantification of the effect of the Trx system onto the import of Tim9 is shown in **Figure 6F** (error bards derived from 3 independent experiments).

**Figure 6.**
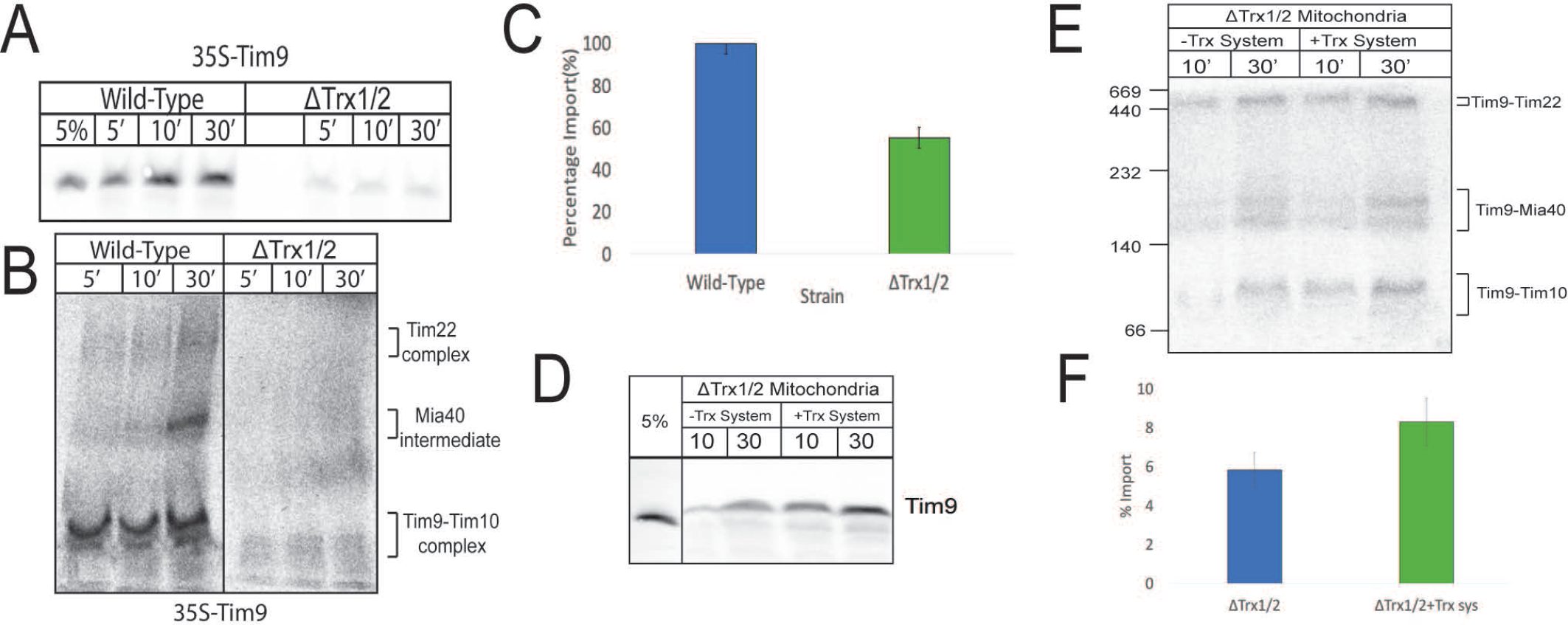
**The Thioredoxin system in the IMS regulates Mia40 substrate biogenesis.** A. Radiolabeled kinetics and import of Tim9 into wild-type and ΔTrx1/2 mitochondria for 5, 10 and 30 minutes. Samples were separated via SDS-PAGE and analysed by autoradiography. 5% in Lane 1 represents 5% of the total translation mix used for import. B. As in A. with the exception that the samples were separated via Blue Native-PAGE. The complexes in which Tim9 participates (Tim9-Tim10 around 70 kDa, Tim as an intermediate with Mia40 around 140 kDa, and Tim9 within the Tim22 complex around 300 kDa) are all indicated C. A histogram quantifying the percentage import of Tim9 after 30 min of import into *Δtrx1/2* mitochondria (green column) relative to the wild type (blue column). Error bars represent standard deviation (n=3). D. Rescue of Tim9 import and assembly in *Δtrx1/2* mitochondria. Ten µg of recombinant pure Trx1 and Trr1 were imported into wild-type or ΔTrx1/2 mitochondria for 5 minutes each with addition of NADPH. The mitochondria were pelleted, washed in breaking buffer, and resuspended in fresh import buffer with radiolabeled Tim9. Samples were taken at 10 and 30 minutes of Tim9 import, separated by SDS-PAGE and analysed by autoradiography. E. As in D with the exception that the samples were separated by BN-PAGE. F. Quantification of Tim9 imports from panel D at a 30 min time point. Error bars represent standard deviation (n=3) (import in *Δtrx1/2* shown in blue; and *Δtrx1/2* supplemented with the full Trx1 system shown in green).

### The Trx in the IMS affects retrograde export of destabilised IMS proteins and protects against mt-ROS induced oxidative protein damage

Given the very broad effects of the Trx system in the cytosol, we wanted to investigate whether the IMS-localised Trx system also has broader effects on mitochondria homeostasis, beyond its direct effects on the MIA import pathway. To address this, we investigated two important pathways where Trx might play a role. First, we asked whether it mediates retrograde export of IMS proteins that are structurally destabilised. Previously, Bragoszewski *et al* (2015) had reported that structurally destabilised proteins of the IMS escape to the cytocol in a retro-translocation pathway, which is facilitated by the chemical reduction of disulfide bonds by DTT, a strong chemical reductant. However, it is still unknown whether a protein system in the IMS may physiologically facilitate this retrograde release pathway. The presence of the Trx system in the IMS that we have shown in this current work prompted us to examine whether the IMS Trx system may have a role in this retrograde release pathway. We used an assay similar to the one described by Bragoszewski *et al* (2015) using the mutant Tim10 C65S as a substrate, since we had previously shown this to be defective in folding in the IMS (Sideris and Tokatlidis 2007; Allen et al, 2003). The results are shown in **Figure 7A** and **7B**. The 35S-labelled Tim10(C65S) was imported into *Δtrx1/2* mitochondria for 10 min, the mitochondria were reisolated by centrifugation and kept for 15 min as follows (i) left untreated (‘-‘ sample, lanes 4 and 5), or (ii) supplemented with the full Trx system (lanes 6 and 7, sequential addition of NADPH/Trx1/Trr1 performed as in **Figure 6**), or (iii) supplemented with DTT (lanes 8 and 9). Samples were then centrifuged to separate the fraction of proteins released from the mitochondria (supernatant, ‘sup’) or still retained within mitochondria (‘M’). Analysis was then done on SDS-PAGE under non-reducing conditions and the Tim10 monomeric fraction that is released from mitochondria in the supernatant was quantified (Figure 7B, three independent experiments). We observe that supplementation of the Trx system in the IMS indeed boosts the retrograde release of Tim10(C65S) outside mitochondria (green column in **Figure 7B**), to a level about half of the total amount that can be released by a harsh chemical treatment with DTT (orange column **Figure 7B**). As a control, the Tim10-Mia40 intermediate that is trapped within mitochondria (indicated with an arrow, lanes 5 and 7) still remains inside the organelle. These data show for the first time that the Trx system in the IMS plays a role in the retro-translocation pathway for structurally destabilised mitochondrial IMS proteins into the cytosol.

**Figure 7.**
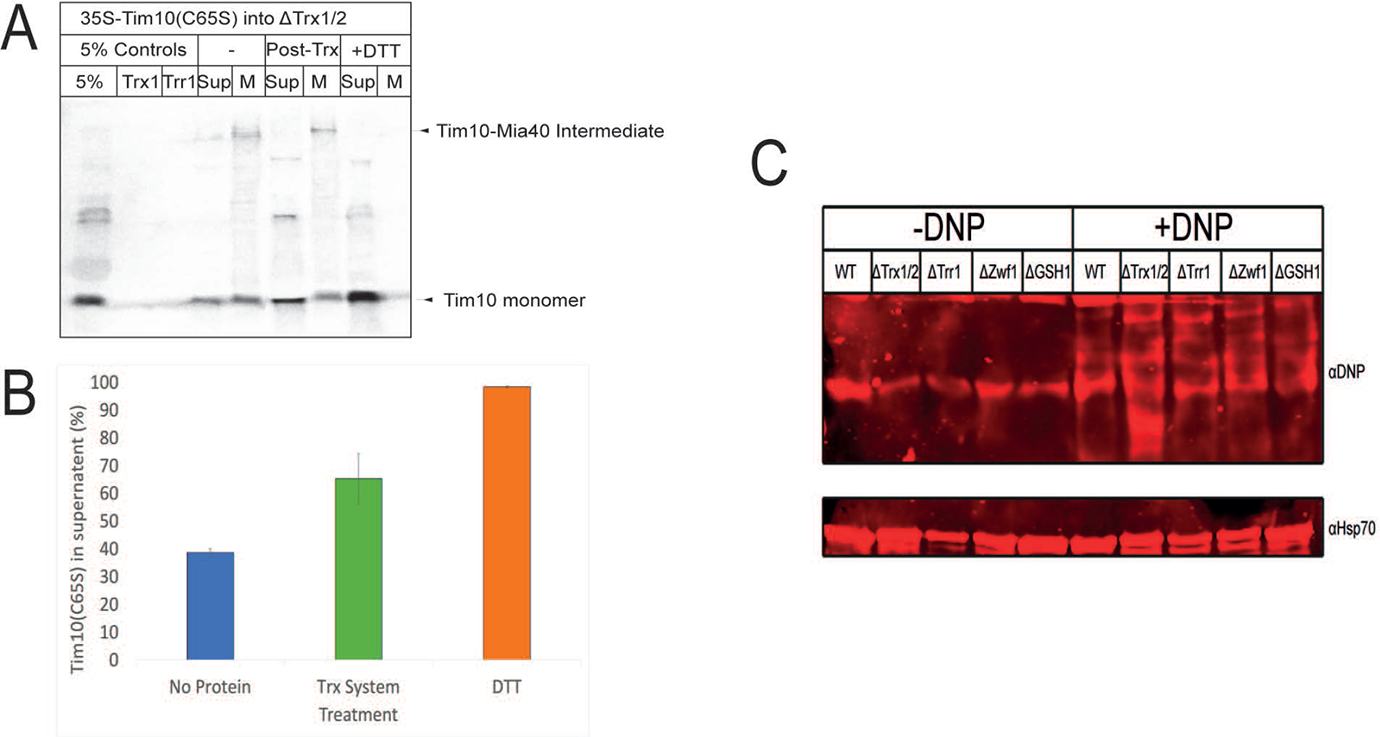
**The IMS Thioredoxin system is implicated in Mia40 substrate retro-translocation and protection against mitochondrial protein carbonylation** A. Release of radiolabeled Tim10(C65S) cysteine trapping mutant from isolated mitochondria. 35S-labelled Tim10(C65S) was imported into wt mitochondria for 30 minutes. The mitochondria were then pelleted, washed, and resuspended in fresh import buffer. 10µg of pure Trx1 and Trr1 were imported for 10 minutes in the presence of NADPH (‘Post-Trx’ samples). As a control, the samples treated with DTT are shown in the last two lanes. The mitochondria were pelleted, the supernatants were TCA precipitated and all samples were separated via SDS-PAGE. B. A histogram quantifying percentage Tim9 in the supernatant (i.e retrograde released out of mitochondria) in A, error bars indicate standard deviation (n=3). Blue column: no protein added to the mitochondria; green column: the full Trx1 system added (‘Post-Trx’ samples in panel A); orange column: DTT added to the mitochondria (‘+DTT’ samples from panel A). C. ‘Oxyblot’ analysis and mitochondrial carbonyl content in wild type, *Δtrx1/2*, *Δtrr1*, *Δzwf1* and *Δgsh1* mitochondria. 50µg of mitochondria were solubilized and protein carbonyls were with either kept in control solution without DNPH (‘-DNP’) or incubated with DNPH (‘+DNP’). Samples were separated by SDS-PAGE and western blotted using an anti-DNP antibody and anti-mtHsp70 as a loading control.

Secondly, we wanted to investigate whether the Trx system abates the generalised oxidative protein damage induced by mt-ROS in the IMS. Protein-bound carbonyls (C=O) represent a marker of global protein oxidation, as they are generated by multiple different reactive oxygen species and consequently levels of protein carbonylation are used as a surrogate for general protein oxidation We used an oxyblot approach whereby mitochondrial extracts from different strains were used to derivatise protein carbonyls of oxidised amino acids with 2,4-dinitrophenylhydrazine, leading to the formation of stable dinitrophenyl (DNP) hydrazone which can then be detected by immunodetection with anti-DNP antibodies (Tsiropoulou and Toyuz 2017). As can be seen in **Figure 7C** the levels of protein carbonylation substantially increase in the Δ*trx1/2* mitochondria (lane 7) compared to all other strains, suggesting a role for the Trx system in controlling the levels of protein oxidation in the mitochondrial IMS. These results point to a protective role for the Trx reductive system against the mt-ROS induced oxidative protein damage in the IMS, potentially providing a compartment-specific shield against irreversible mitochondria damage.

## Discussion

The reductive Trx system is key to the antioxidant response of cells and is widely conserved across species (Draculic *et al*, 2000, Garrido & Grant, 2002). The cytosolic isoforms of the Trx system, Trx1 and Trr1, were found to be present in the mitochondrial IMS of *Saccharomyces cerevisiae* (Vögtle *et al*, 2012). This raised the question whether the Trx system itself might be an as yet missing machinery in the mitochondrial IMS, equivalent to other thiol-reducing protein systems that maintain a critical balance between an oxidising and a reducing pathway in compartments where oxidative folding occurs, i.e. the bacterial periplasm and the ER lumen (Manganas *et al*, 2017; Cardenas-Rodriguez & Tokatlidis, 2017). In the bacterial periplasm, the mitochondrial ancestor, the reducing DsbC/DsbD proteins accepts electrons from the cytosolic thioredoxin. In this bacterial compartment, the oxidative and reducing mechanisms are kept kinetically separated by steric impediment between the oxidative pair (DsbA/DsbB) and the reducing pair (DsbC/DsbD) (Rozhkova *et al*, 2004). In the ER, the reducing power comes from both GSH and NADPH, and like in bacteria, the cytosolic thioredoxin system (Poet et al EMBO J 2017). Unlike bacteria though, both the oxidative and reductive parts of oxidative folding are ensured by the same PDI protein that can either oxidise or reduce disulfides (Ellgaard *et al*, 2018, Oka *et al,* 2017). The redox balance in the ER is determined by a feedback regulation of PDI by the oxidoreductase Ero1, with the oxidised/reduced ratio of this protein determining a reducing or oxidising flux (Sevier *et al*, 2007).

We addressed here whether the cytosolic thioredoxin system can serve as the direct reducing machinery in the mitochondrial IMS, as it is located in this compartment as well via a dual localisation mechanism. This would represent a key difference to the bacterial and ER compartments where the cytosolic thioredoxin system is physically segregated from these compartments. We first confirmed that both Trx1 and Trr1 can be imported into mitochondria and co-localise to this compartment (**Figure 1B** and **Figure 1C**). We have also seen that Trx2 (which is almost identical to Trx1) also gets imported to the IMS, but we focused on the import of Trx1 for the bulk of this work. Interestingly, the import of Trx1 and Trr1 does not depend on ATP or Δψ (**Figure 1C**), which are the major energy requirements for the import of the majority of proteins to the mitochondrial matrix or the mitochondrial inner membrane by the TIM23 and TIM22 complexes (Geissler *et al*, 2000, Pfanner & Neupert, 1985, Pfanner & Neupert, 1987, Pfanner *et al*, 1987).

The main import pathway into the mitochondrial IMS is the MIA pathway. However, the import of both Trx1 and Trr1 is independent of the main MIA pathway components Mia40 and Erv1 (**Figure 1D**). This independence from the MIA import pathway is an emerging common theme for other proteins that are involved in regulation of the oxidative folding pathway. Such examples are the thiol peroxidase Gpx3 (Kritsiligkou *et al*, 2017) and the final electron acceptors for the pathway like cytochrome c and cytochrome C peroxidase Ccp1 (Dabir *et al,* 2007; Allen *et al,* 2005). Gpx3 import is boosted by an N-terminal extension produced by alternative cytosolic translation upon hydrogen peroxide stress (Kritsiligkou *et al*, 2017), whilst Cytochrome C is imported via an unusual Tom40-dependent pathway (Mayer *et al*, 1995). The import of Trx1 (and Trx2) and Trr1 is yet distinct, and does not follow any of the known pathways, while it is independent of the cysteines present in the proteins (**Figure 1E**). Additionally, the presence of either Trx1 or Trr1 in the IMS was also not required for the import process of these proteins (**Figure 1F**). This is in clear contrast to other proteins dually localised in the IMS and the cytosol, like the SOD1 chaperone CCS1 and SOD1. These proteins interact with each other in both compartments and targeting of SOD1 in the IMS is determined by the prior presence of CCS1 in the IMS (Sturtz *et al*, 2001). It will therefore be interesting to tease out the mechanism of the yet unknown pathway that ensures import of Trx1 and Trr1 in the IMS independently of each other.

The concurrent presence of Trr1 with Trx1 in the IMS fraction ensures the functionality of the Trx reductive system in this compartment. The dependence of reduction of Trx1 on Trr1 supports the concept that the IMS-localised system follows the same mechanism as in the cytosol (**Figure 2A**), but clearly with a different set of substrate proteins which are localised in the IMS. We found that Mia40 interacts with Trx1 *in vitro* and *in organello* **(Figure 2B** and **Figure 2C**) and that it can be reduced by the Trx system (**Figure 3**), which illustrates Mia40 as one of the substrates of Trx1 in the IMS. Given that oxidative folding requires a balance between an oxidative and a reductive reaction (Banci *et al*, 2009, Grumbt *et al*, 2007), ensuring a Trx-dependent reductive reaction in parallel to the Mia40-dependent oxidative reaction is crucial for optimal mitochondria biogenesis and cellular physiology.

NADPH provides the crucial electron pool for the function of both the Trx and Grx systems. The cytosolic population of NADPH is produced by reduction of NADP concomitantly with the oxidation of glucose-6-phosphate (G6P) to 6-phosphoglucolactone (6PGL), the first step of the pentose phosphate pathway, which is catalysed by G6PD (Nogae & Johnston, 1990), The deficiency of this enzyme is the most common enzyme deficiency in humans (Minucci *et al*, 2012). Accordingly, G6PD is involved in the adaptive response to oxidative stress in all organisms including *S. cerevisiae* (Izawa *et al*, 1998) and serves as the key supplier of NADPH for the Trx reductive function. Given the dual localisation of the Trx system in the IMS, is a separate pool of NADPH present in the IMS or is the cytosolic NADPH communicated across the OM to the IMS? To address this, we tested whether a null mutant in the *zwf1* gene in *S. cerevisiae* (the yeast homologue of human G6PD) that lacks G6PD activity in the cytosol showed any IMS-level defects. Indeed, this was the case as we find that the MIA machinery (and no other import components) is substantially and specifically affected due to a dysregulation of its redox balance, remaining in a mostly oxidised state in a *zwf1*Δ strain (**Figure 4A and B**). This is important because it was previously shown that the functional Mia40 pool is in a mixed reduced/oxidised state in both yeast and human cells (Kojer *et al,* 2012, Erdogan *et al*, 2018). A Mia40 redox imbalance would conceivably affect the optimal function of mitochondria. Although it can be argued that the cytosolic NADPH can reach the IMS through the OM porin channels to power the Trx reductive system in this compartment, this is not yet known and would be important to address in future studies. No matter what the exact mechanism of NADPH transport across the OM, we show here for the first time that a crucial mitochondrial import pathway, the MIA pathway, is subject to cytosolic metabolic control. This communication becomes critical because of the presence in the IMS of the thioredoxin system that depends on NADPH to operate. The import of Trx1 and Trr1 in the IMS of ΔG6PD mitochondria, combined with the addition of NADPH to provide the reducing power for the Trr1/Trx1 system restored the redox state of Mia40 to wt levels and the import of MIA substrates (**Figures 5-6**).

On the other hand, the glutaredoxin system was also proposed to provide a reductive force in the mitochondrial IMS. However, a KO yeast for glutaredoxin reductase (Glr1) did not affect the redox state of Mia40 (Kojer *et al,* 2012). In particular, the cytosolic isoform of the Grx system, Grx2, was proposed to be an active reductive component in the IMS (Kojer and Riemer, 2014). However, unlike the results presented here where Trx1 and Trr1 can be imported into mitochondria, no influence of exogenous Grx2 was seen when presented to mitochondria expressing an IMS-localised roGFP2 probe that responds to redox changes in the IMS. Thus, we conclude that the Trx system is likely a major reductive machinery in the mitochondrial IMS, although likely not the only one. We propose that the Trx system regulates the import capacity of Mia40 in the IMS by maintaining a functional reductive/oxidised balance (**Figures 5-6**).

It is intriguing that the interaction of Mia40 and Trx1 proceeds independently of the active site hydrophobic cleft of Mia40, which is required for the interaction of Mia40 both with its substrates and with Erv1 (Banci *et al,* 2011) . This suggests that the interaction Mia40 follows an interaction mechanism with Trx which is novel and based on structurally distinct requirements. The electrochemical potential of Mia40 is -290mV and that of the CX3C and CX9C motifs of its import substrates is -310mV and -390mV, correspondingly (Riemer *et al*, 2011; Banci *et al*, 2009). This suggests a thermodynamic impediment for a reducing (isomerisation) role of Mia40 towards its classic substrates. The same situation applies for a reaction between Mia40 and Trx1 (E°= -230 to -280 mV). However, all the E° were determined at pH 7 and controlled conditions. The environment within the IMS might allow these reducing reactions to take place, due to the dynamic response of this compartment in changes in the cytosol (Kojer *et al,* 2012), which may change the exposure of the reactive sites of the proteins involved, favouring the reducing reactions to happen.

The dual localisation of the Trx system in the mitochondrial IMS in addition to the cytosol is likely not an oddity of yeast cells but also true for human cells. In support of this, Nakao *et al* (2015) found human Mia40 among a specific set of proteins that were pulled down by a human thioredoxin thiol-trapping mutant (similar to the one we used here for yeast Trx1) in an unbiased search for protein substrates of the human Trx protein. Our study provides a mechanistic/functional framework for such an interaction and highlights for the first time the Trx machinery as potentially the reductive arm to the oxidative folding process in the IMS.

Taken together our data highlight the dual localisation of the cytosolic thioredoxin system in the mitochondrial IMS as a mechanism to maintain optimal mitochondrial fitness. We have shown that this role may be served in at least three different ways. Primarily, by maintaining the redox balance of Mia40 and supporting the efficient import of its substrates. Additionally, the Trx system seems to assist the efficient clearance of structurally destabilised proteins (containing non-native disulphide bonds) via a retrograde export to the cytosol that allows their clearance by the cytosolic proteasome system. Although the details of the function of the Trx system in this retrograde escape process and the extent of proteins that may affected by it remain to be clarified, the discovery of the IMS Trx system as the first protein machinery involved in this process is an important step.

Finally, the Trx system provides a general protection against mtROS-induced oxidative damage of IMS proteins, as revealed by our mitochondrial protein carbonylation analysis. Such a role might have links to another antioxidant protein that is also dually targeted in the IMS, the thiol peroxidase Gpx3 (Kritsiligkou *et al*, 2017), which is not only a critical partner for Mia40 but also is a key player in the protection against lipid peroxides.

Collating all our data a working model for the presence of the Trx machinery in the IMS is depicted in **Figure 8**.

**Figure 8.**
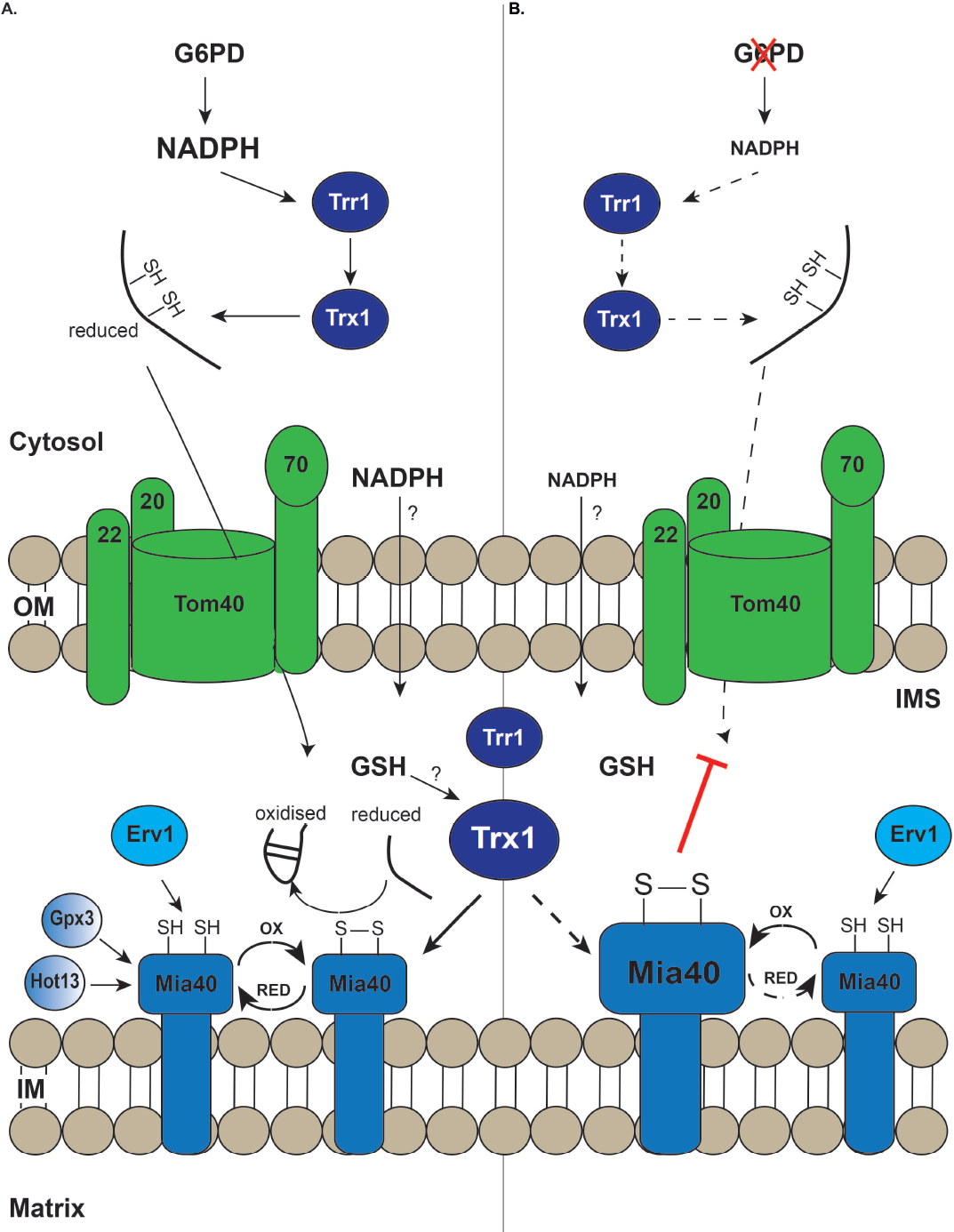
**Proposed model for the Trx machinery redox regulation in the mitochondrial IMS** A. Under normal conditions cytosolic NADPH feeds with electrons the Trx system in the cytosol to keep MIA substrates in a reduced state to allow them to translocate through the Tom40 channel. In the IMS, the substrate is oxidised by Mia40. Reduced Mia40 is recycled back to its oxidised form by Erv1 with the help of Gpx3 and probably Hot13. Here, Mia40 is present in a mixed reduced and oxidised state, where the Trx system is responsible of keeping Mia40 in a reduced state. NADPH delivered from the cytosol into the IMS (and therefore the function of G6PD to generate NADPH) is crucial to feed electrons to the IMS-localised Trr1 and Trx1 and keep them in a functional state. In this way, the Trx protein machinery in the IMS functions as the reductive part of the oxidative folding pathway (in which GSH may also contribute to maintain a redox balance). B. When G6PD is absent the Trx system in the IMS is blocked disturbing the redox balance of Mia40 which is now mainly in an oxidised form. Disruption of the redox balance of Mia40 affects the import, folding and assembly of Mia40 substrates. In addition to its role in the oxidative folding pathway, the Trx machinery in the IMS serves additional functions in the retro-translocation of structurally destabilised proteins back to the cytosol for proteasomal clearance and as a protective system against ROS-induced oxidative damage in mitochondria.

The thioredoxin system therefore represents an important component of protein biogenesis and quality control in the mitochondrial IMS, with critical roles for optimal mitochondria health and response to stress.

## Acknowledgements

We thank members of our lab for discussions and comments on the manuscript, Donna McGow (Institute of Molecular Cell and Systems Biology, University of Glasgow) and Nitsa Katrakili (Institute of Molecular Biology and Biotechnology, Foundation for Research and Technology, Heraklion, Crete, Greece) for expert technical assistance in preliminary phases of this work. We thanks Prof Chris Grant (University of Manchester for the gift of strain *Δtrx1/trx2*. MC-R was supported by a PhD studentship from CONACyT (Consejo Nacional de Ciencia y Tecnologia Mexico) and RE by a Lord Kelvin-Adam Smith PhD studentship. Work in our laboratory was supported by strategic funds of the University of Glasgow, the Scottish Universities Life Science Alliance and the Scottish Funding Council (HR07019), the Royal Society (Wolfson research merit award to KT Grant Code: WM120111), the Wellcome Trust (Grant Code: 202308/Z16/Z), EU COST action ‘EU-ROS’ (BM1203), EU COST Action ‘FeSBioNet’ (CA15133) and UKRI-BBSRC (grants BB/R009031/1 and BB/T003804/1).

## Author contributions

MCR, PM and RE performed experiments, made figures, analysed data and wrote the manuscript with KT. EK and AC performed experiments and analysed data. EL purified proteins. KT designed the study, secured funding, directed the research, analysed data and wrote the manuscript with MCR and PM and input from all authors.

## Materials and Methods

### Yeast and growth conditions

The yeast strains used are derivatives of the BY4741 (MATa, his3Δ1, leu2Δ0, met15Δ0, ura3Δ0). The yeast strains *Δzwf1*, *Δtrx1*, *Δtrr1* and *Δgsh1* were purchased from GE Dharmacon (Lafayette, CO, USA). Yeast cells were grown in rich (YPD or YPlac) media at 30°C. The media contained 2% (w/v) carbon source (glucose for YPD and lactic acid for YPLac), 1% (w/v) yeast extract and 2% (w/v) peptone. In the case of YPlac, the pH was adjusted to pH 5.5 to ensure conversion of the acid species to lactate. Additionally, the media was supplemented with 76 mg/L methionine for the growth curve, growth spot tests and isolation of mitochondria from *Δzwf1* and *Δtrx1/2* yeast.

### Purification of mitochondria

Wild-type and mutant mitochondria were isolated from the respective yeast strains were isolated as previously described (Glick, 1991; Banci *et al*, 2009). The yeast strains were grown at 30°C in medium containing 1% (w/v) yeast extract, 2% (w/v) bacto-peptone, and 2% lactate (v/v) pH adjusted to 5.5 (YPLac).

### Protein import into purified yeast mitochondria

Mitochondrial ^35^S-labeled precursor proteins were synthesized using the TNT SP6-coupled transcription/translation kit (Promega) and plasmid-vectors pSP64 containing the genes of interest. Additionally, 6XHis tagged-Trx1 and Trr1 recombinant proteins were cloned in PET24 vectors and purified by column affinity onto Ni-NTA beads (Qiagen). The precursor was imported in 50 *μ*g of the indicated yeast mitochondria in the presence of 2 mM ATP and 2.5 mM NADH at 30°C. Mitochondria were resuspended in 1.2 M sorbitol and 20 mM HEPES (pH 7.4), followed by a treatment with 0.1 mg/mL proteinase K (PK) to remove unimported material for 20 min on ice and PK inactivation for an extra 10 min on ice in the presence of 2 mM PMSF. For the fractionation experiments, the mitochondria were treated with 0.1mg/ml trypsin for 30min on ice to remove the unimported material, followed by inactivation with 1mg/ml SBTI (soybean trypsin inhibitor) for 10min on ice. The mitochondrial pellets were washed twice with 1.2M sorbitol and 20mM HEPES (pH 7.4) to ensure the removal of the unimported material, as well as the trypsin that was added, before proceeding with the fractionation. Mitoplasts were produced by resuspending mitochondria in import buffer (Sideris & Tokatlidis, 2010) at a concentration of 5 mg/mL and diluting 10 times with 20 mM HEPES (pH 7.4) in the presence or absence of 0.15 mg/mL PK for 30 min on ice. The supernatant was kept for trichloroacetic acid (TCA) precipitation (Chatzi *et al*, 2013). For carbonate extraction, isolated mitochondria were resuspended in ice-cold, freshly prepared 0.1 M Na_2_CO_3_ and incubated on ice for 30 min; the pellet was then recovered by centrifugation (55000g, 30 min, 4°C). Finally, samples were resuspended in Laemmli sample buffer with β-mercaptoethanol as indicated, analysed by SDS-PAGE and visualized by immunodetection or autoradiography. For the import of proteins into mitochondria with disrupted membrane potential (Δψ), NADH was omitted from the import mix. Instead, valinomycin, a membrane potential disrupting chemical, was added at final concentration of 1mM. For the import of proteins into mitochondria depleted of ATP, ATP was omitted from the import mix. Instead, 10μM oligomycin and 10mU/μl apyrase were added to the import mix and were incubated with the mitochondria for 10min prior to the addition of the precursor.

### Trapping of mixed disulfide intermediates after import of proteins

For trapping mixed disulfide intermediates during the import process, the import assay was performed as described previously and the redox state of the proteins in the reaction was arrested by adding 20mM of N-ethylmaleimide (NEM) for 5min on ice and the samples were analysed on SDS-PAGE in the absence of a reducing agent.

### Mitochondria retro-translocation release assay

The assay was performed essentially as described by Bragoszewski *et al* (2015), but using the Tom10(C65S) as a structurally destabilised IMS protein. Briefly, 35S-labelled Tim10(C65S) was imported into wt mitochondria for 30 minutes. The mitochondria were then pelleted, washed, and resuspended in fresh import buffer. One aliquote was left untreated. In another aliquot, 10µg of pure Trx1 and Trr1 were imported for 10 minutes. The mitochondria were pelleted, and supernatant was collected as the released retro-translocated protein, keeping the pellet as the protein still retained within mitochondria. The supernatants were precipitated with ice-cold 10% trichloroacetic acid and all samples were separated via SDS-PAGE and analysed by autoradiography.

### Alkylation shift assay

The alkylation shift assay was performed using either AMS (Sigma) or mal-PEG 5000 (Sigma). First, 100µg of mitochondria were precipitated with 10% final concentration ice cold TCA. The protein pellet was resuspended in 8µl 15mM final concentration of AMS in TES (50mM Tris-HCl pH 8, 3mM EDTA and 3% SDS) or mal-PEG in HES buffer (50mM HEPES-KOH pH 7.4, 10mM EDTA and 3% SDS) or only with TES/HES buffer and cover from light with aluminium foil. Samples were incubated at 30°C for 30min and then at 37°C for 30min more. Finally, 17µl of 4X Laemmli sample buffer were added to the reaction.

### ***In vitro* interaction assays**

All proteins (50*μ*g) used were precipitated with saturated ammonium sulfate and then resuspended in an appropriate buffer as indicated. For the control samples (usually lanes 1-4 in each image as indicated), they were then treated with either 200μM of freshly prepared H_2_O_2_ (for oxidation. shown as ‘ox’) or with 20mM DTT (for reduction, shown as ‘red’) for 30min. Where required, 5mM NADPH was added to Trr1 for 30min. Each reaction was performed in 1ml of 50mM Tris-HCl pH8 buffer at 30°C for the indicated time followed by TCA precipitation. The samples were then treated with 15mM (final concentration) of the alkylation agent AMS in TES buffer with the same conditions as described above.

### Chemical crosslinking

The interaction between Mia40 and either Trx1 or Trx2 was tested by crosslinking with glutaraldehyde (GA). Equimolar concentrations of purified Trx1His, Trx2His or Mia40Δ290 His in 50 mM Hepes-KOH pH 7.4 buffer were incubated either alone or in the presence of GA at a final GA concentration of 0.1 %, on ice, and for varying times between 20 seconds and 10 minutes. The crosslinking reaction was stopped by addition of 0.1 M ice-cold TrisHCl pH 8.0. Samples were then analysed by SDS-PAGE and immunoblotting using anti-His rabbit polyclonal antibodies.

### Isothermal titration calorimetry

Isothermal titration calorimetry (ITC) was performed essentially according to Sideris *et al* 2009. We used in the syringe 1 mM Mia40Δ290 His (Sideris *et al*, 2009; Mia40 lacking the first 290 amino acids that are not involved in its function) and 0.1 mM TrxC30/33SHis in the cell. The dissociation constant Kd was calculated using the Origin software and found to be approx. 30 μM.

### Oxyblot mitochondrial protein carbonylation detection

Our analysis was based mainly on the guidelines of the Abcam oxidised protein western blot detection kit (ab178020). Briefly, 50μg of wild-type, Δ*trx1/2*, Δ*trr1*, Δ*zwf1* and Δ*gsh1* mitochondria were solubilised in 6% SDS solution +0.5% Triton X-100 and incubated at RT for 10 minutes. The carbonyl reactive compound, 1X DNPH or the derivatisation control solution was added to the solubilised mitochondria and incubated at RT for 15 minutes. The free DNPH was quenched with neutralisation solution, and 6X Laemmli buffer was added gto the samples, which were separated by SDS-PAGE followed by western blotting. The anti-DNPH antibody was used to indirectly detect the carbonyl products via the DNPH moiety conjugated to them.

---

**Supplementary Figure 1.**
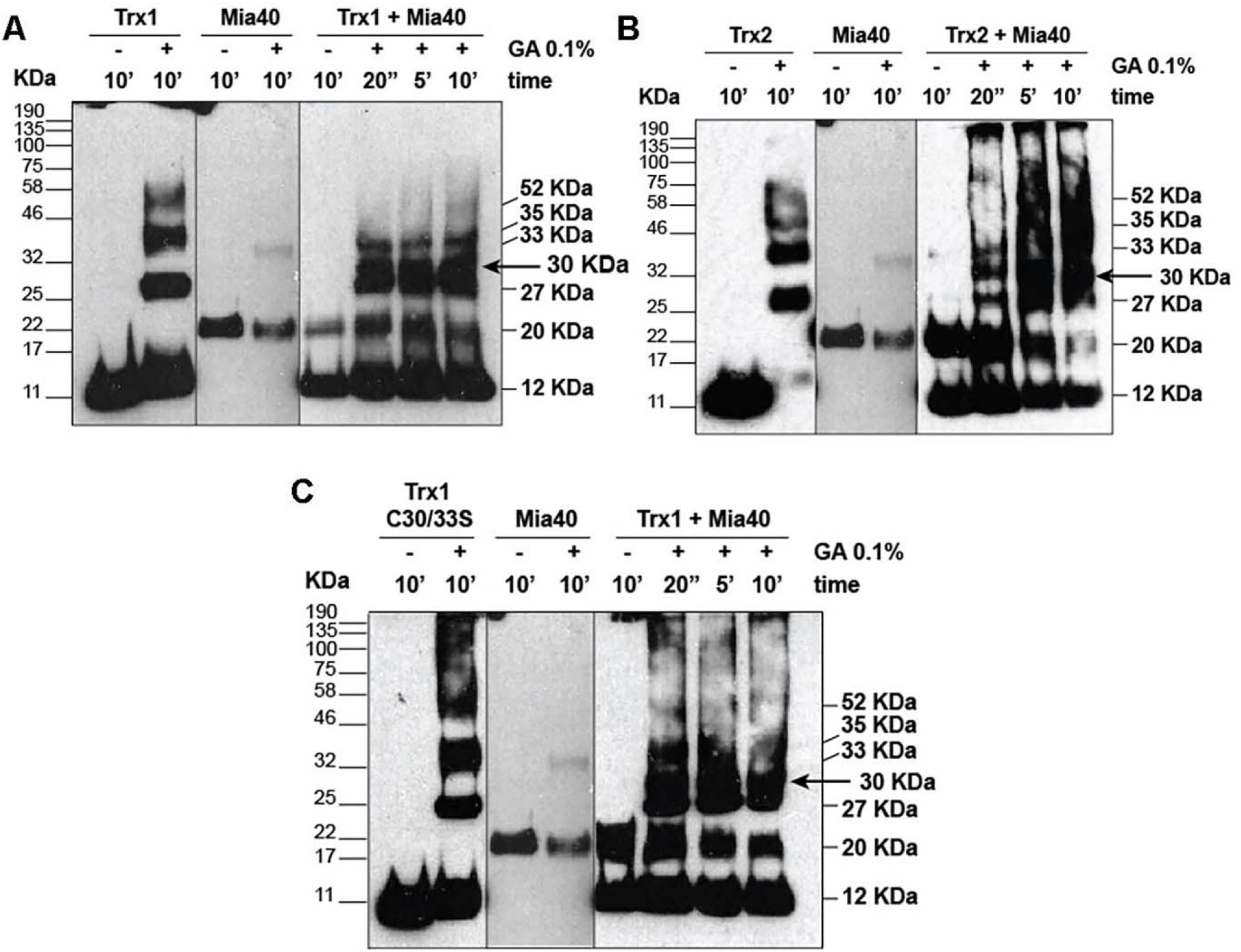
Chemical crosslinking between Trx1 or Trx2 and Mia40 shows they interact *in vitro*. Gluteraldehyde was added at 0.1 % final concentration and incubated for the indicated times (between 20 seconds to 10 minutes). After crosslinking and SDS-PAGE protein adducts were revealed by anti-His antibodies (both Trx1, Trx2 and Mia40 proteins were purified as 6His-tagged proteins). The additional band that is generated only when either Trx1 or Trx2 and Mia40 are incubated together is shown with an arrow. Approximate mol. Weights of the crosslinked products are indicated to the right of the gels, and the molecular weight markers to the left. A. Trx1 and Mia40 interaction, B. Trx2 and Mia40 interaction, C. Trx1C30/33S double mutant interaction.

